# Cellular mechanisms underlying social regulation of the posterior tubercular nucleus in zebrafish (*Danio rerio*)

**DOI:** 10.64898/2026.01.27.701991

**Authors:** Carrie L Adams, Emily M Scott, Fadi A Issa

## Abstract

Social status profoundly influences animal behavior through neural plasticity, yet the cellular mechanisms that mediate reconfiguration of neuromodulatory systems remain poorly understood. Here, we investigated status-dependent structural changes in the posterior tubercular nucleus (PTN) of adult zebrafish. Animals were assigned to four social conditions: communal, isolated, dominant, or subordinate. Using markers for cell proliferation (PCNA) and birth-dating (BrdU), we demonstrate that social dominance significantly enhances cell proliferation, leading to an increased population of PTN dopaminergic neurons. In contrast, subordinate and isolated fish exhibited suppressed proliferation and elevated expression of superoxide dismutase 1 (SOD1), suggesting that chronic social stress induces an oxidative burden that may lead to neuronal loss. Furthermore, we identified evidence of neurotransmitter phenotypic plasticity; subordinate fish displayed a significantly higher ratio of glutamatergic (*vglut2a*) to dopaminergic (*dat*) expression in PTN neurons compared to dominants, suggesting a status-dependent shift in neuromodulatory identity. Multivariate principal component analysis showed distinct neurobiological profiles that separate social ranks, suggesting that status-dependent plasticity is a coordinated multi-modal response whereby increased BrdU and PCNA expression clustered with the dominant profile while increased expression of cellular stress and shift to glutamate cellular identity clustered with social subordinate and isolate profiles. Collectively, our results improve our understanding of how social experience reshapes the zebrafish brain through integrated changes in cell proliferation, cellular shift in neurotransmitter identity and regulation of cellular viability; thus, providing a potential mechanism for the maintenance of stable behavioral phenotypes in competitive social environments.

## INTRODUCTION

Social aggression is a fundamental adaptive behavior observed across diverse animal taxa, serving as the primary mechanism for the resolution of resource competition. In many species, agonistic interactions do not merely result in transient conflict but lead to the establishment of stable dominance hierarchies that optimize the distribution of important resources (Nelson and Trainor, 2007; Oliveira et al., 2011; Flanigan and Russo, 2019). These hierarchies provide a predictable social structure that reduces the fitness costs associated with aggression (Drummond et al., 2002; Herberholz et al., 2007; Clements et al., 2023a). However, maintaining a specific social rank requires significant physiological and neurological plasticity. While the ecological drivers of dominance are well-defined, our understanding of the neurobiological substrates that allow animals to transform social experience into a long-term behavioral phenotype remain poorly understood.

Zebrafish have emerged as a model organism for investigating the intersection of social behavior and neural plasticity (Teles et al., 2016; Clements et al., 2018; Carver et al., 2021; Dunlap et al., 2021, 2021). Their utility is derived from a combination of high genetic tractability and rapid development (Kalueff et al., 2014; Stewart et al., 2014). Critically, zebrafish exhibit a sophisticated social repertoire characterized by aggressive social interactions that lead to the establishment of stable dominance relationships in paired or group settings. These status-dependent relationships are defined by distinct behavioral archetypes: dominant individuals exhibit proactive territorial aggression and increased locomotor activity, whereas subordinates display reactive avoidance, social withdrawal, and a heightened sensitivity to environmental threats (Miller et al., 2017; Nakajo et al., 2020). Such behavioral divergence suggests that social rank is encoded through the plasticity of highly conserved circuits that integrate social cues with motor output.

The divergence in behavior following rank establishment is largely modulated by the “Social Decision-Making Network” (SDMN). The SDMN comprises the social behavior network and the mesolimbic reward system is evolutionarily conserved across many vertebrates (O’Connell and Hofmann, 2012). This conservation suggests that vertebrate social diversity arises from subtle variations on a fundamental neural theme, where internal physiological cues are integrated with external stimuli to produce adaptive behavioral responses. Central to the SDMN is the dopaminergic (DA) system, which modulates motivation, arousal, and motor control (Schweitzer & Driever, 2009). In the teleost brain, the Posterior Tubercular Nucleus (PTN) represents a critical diencephalic DA center and is considered a functional homolog to the mammalian A11 group (Rink & Wullimann, 2002). Characterization of the DA system in teleost has identified that PTN DA neurons express specific dopaminergic receptor subtypes that are essential for evaluating the salience of environmental stimuli (O’Connell et al., 2010). Furthermore, neurochemical profiling indicates that teleost DA neurons share conserved molecular markers with their mammalian counterparts, confirming that the PTN is an integral component of the vertebrate reward and decision-making apparatus (O’Connell et al., 2013). This unique neuroanatomical position allows the PTN to integrate and relay social cues to drive specific motor patterns required for either swimming or startle-escape behaviors (Clements et al., 2023).

The role of dopamine in social regulation is multifaceted, specifically in how it modulates an organism’s responsiveness to social feedback. DA signaling within the diencephalon helps categorize interactions as either rewarding or aversive, thereby guiding future decision-making (Antunes et al., 2022). Consequently, the dopaminergic system is highly sensitive to the chronic stress of subordination. In zebrafish and other fish species, chronic social defeat leads to a downregulation of brain DA levels, resulting in a shift toward energy-conserving, reactive behavioral strategies (Miller et al., 2017; Antunes et al., 2022). In contrast, dominant status is typically supported by robust DA activity that facilitates proactive exploration and persistence. While these neurochemical shifts are established, the cellular remodeling within the PTN that facilitates these long-term behavioral changes is only beginning to be elucidated.

A recent report demonstrated that the PTN undergoes significant morphological remodeling in response to sustained social experience (Heagy et al., 2024). Morphometric analyses revealed that dominant zebrafish have significantly higher number of DA neurons in the PTN compared to subordinate (Heagy et al., 2024). Notably, this increase in cell number is inversely correlated with synaptic density, suggesting a complex interplay between cell proliferation and synaptic pruning during the formation of social dominance. A previous longitudinal study indicates that these structural differences are not innate but emerge between 7 and 14 days following dominance formation and could be reversed if subordinates were provided the opportunity for social rise or whether dominants were forced into social submission (Heagy et al., 2024). This suggests that the PTN is a site of socially induced cellular plasticity, where the environment sculpts the nucleus to match the animal’s ecological niche within the hierarchy. However, an outstanding question remained unanswered: what are the cellular mechanisms that underly this shift in plasticity?

Here, we sought to understand the mechanisms underlying the morphological changes within the PTN that occur as a direct consequence of social status. Specifically, we tested the hypothesis that social rise will promote cell proliferation, while social stress will lead to reduction in cellular viability and potential loss. Our results show that status-dependent cellular plasticity in PTN neuronal number is mediated by multiple processes consisting of cell proliferation, a shift in neurotransmitter identity, and cellular oxidative stress working synergistically to regulate the morphological architecture of the PTN. By defining the mechanisms of PTN plasticity within the context of a conserved vertebrate SDMN, our results improve our understanding of how social experience remodels the brain to sustain adaptive behavioral strategies in fluctuating social environments.

## METHODS & MATERIALS

### Animal husbandry

Zebrafish transgenic line Tg*(dat:*egfp) and double transgenic line Tg*(dat:*egfp), Tg(*vglut2a:*rfp) were raised and maintained in grouped-housed tanks consisting of 10-20 mixed sex adult zebrafish at 28 °C, pH of 7.1-7.2, 10dark/14light cycle, and fed live brine shrimp and pellet food twice daily. Experiments were approved by East Carolina University’s Institutional Animal Care and Use Committee.

### Isolation, pairing, and behavioral observations

Adult male Tg(*dat*:egfp) zebrafish, ranging from 3-6 months old, were isolated physically and visually from conspecifics for one week in 23×13×6 cm individual tanks, to minimize prior social experience. Following isolation, two male zebrafish of equal size, age, and identical genetic background were paired continuously for two weeks. Aggressive interactions were observed daily for 2-3 minutes at mid-day, and the number of attacks and retreats was quantified as described elsewhere (Miller et al., 2017). Social dominance was determined daily for each pair based on the total number of aggressive attacks (biting and chasing) and defensive retreats. Fish with the most attacks were considered dominants, while those that retreated the most were considered subordinates. While female zebrafish exhibit agonistic behaviors and establish short-term dominance hierarchies, their diurnal ovulatory cycles and associated hormonal fluctuations result in inconsistent aggression and transient social structures (Teles and Oliveira, 2016). Given that the objective of the study was to investigate the long-term effects of stable social dominance on the mechanisms underlying morphological plasticity in the PTN, females were excluded. This ensured a more controlled experimental framework, as male zebrafish maintain significantly more stable dominance relationships than their female counterparts.

### BrdU Injections

Adult male Tg(*dat*:egfp) zebrafish of known social status were injected with BrdU at two mid-points during the pairing (days 8 and 10). We selected these two time points because a prior report demonstrated that the number of PTN cells between dominants and subordinates begins to differ starting at 9-days of social interactions (Heagy et al., 2024). Communals were injected and returned to their tanks for 48 hours to recover. Isolates were injected on days 2 and 4 of isolation. Bromodeoxyuridine (BrdU) (G3G4-BrdU-s) was administered intraperitoneally. A stock solution of BrdU (10 mM) was prepared in sterile phosphate-buffered saline (PBS), filtered through a 0.22 μm filter, and stored at -20°C until use. On the day of injection, the solution was diluted to 2.5 mg/mL in PBS. Adult zebrafish were anesthetized using 0.025% tricaine methanesulfonate (MS-222) and placed ventral side up on a damp sponge. A capillary glass filament needle with 50 μL of BrdU solution was injected into the intraperitoneal cavity, posterior to the pectoral fins, at an approximate angle of 30 degrees (Tea et al., 2019). Fish were then transferred to recovery tanks and monitored for normal swimming behavior for 2-3 hours, then returned to their appropriate tanks to interact with their paired counterpart.

### Euthanasia

Animals were euthanized in a heavy dose of tricaine (0.25%) for 15 minutes followed by sudden emersion in iced water for 15 minutes prior to brain extraction.

### Histology

Extraction of zebrafish brains occurred on days 8 (PCNA) and 14 (BrdU, SOD1, and Neurotransmitter shifts) and placed at 4 °C overnight in a 2 mL microcentrifuge tube containing 1 mL of 4% paraformaldehyde. Brains were washed 4-5 times for 25 minutes in 1x PBS. After washing, brain tissues were incubated overnight at 4 °C in 15% sucrose for cryoprotection. The next day, brains were transferred to a 30% sucrose solution and incubated overnight at 4 °C. Brains were then placed in a mold filled with OCT, oriented for sagittal sectioning, and flash-frozen in liquid nitrogen. OCT-embedded brains were placed in a cryostat (Microm HM 550) at - 23°C and sliced at 45 µm and mounted onto Superfrost Plus Microscope slides (Fisher Scientific).

### BrdU staining

BrdU was used as a marker of cell proliferation. Slides were kept face up throughout the staining protocol. Following the Gilmour protocol, brain slices were washed 3-4 times for 5 minutes each in PBS, then incubated in a 2mM HCl solution at 37°C for 30 minutes. After incubation, the slides were washed 2-3 times using a 0.1 M borate buffer at pH 9.0 for 5 minutes each. Brain slices were washed 2-3 times for 5 minutes each in PBST, then placed in a 5% bovine serum and 2% goat serum blocking buffer for 2 hours following washes. Another series of PBS washes (3-4 times for 5 minutes) was performed, and slices were stained with a mouse polyclonal anti-BrdU primary antibody at a concentration of 1:300 (Tea et al., 2019). All slides are appropriately placed overnight in the fridge at 4 °C. Following primary antibody incubation, an additional 3-4 PBS washes were performed, and slices were incubated for 3 hours at room temperature in goat anti-mouse Alexa Fluor-555 secondary antibodies (Thermo Fisher, REF: A21422, Lot: 2978487) diluted at a 1:750 concentration. Secondary antibodies were washed in PBS, and the slides were mounted with one drop of Permount SP15-500 toluene solution (Thermo Fisher). Slides were cured in the fridge for 24 hours, then sealed with clear nail polish for protection.

### PCNA and SOD1 staining

Proliferating Cell Nuclear Antigen (PCNA) was used as a marker of cell proliferation. Brain slices were washed 3-4 times for 5 minutes each in PBS, then placed in 5% bovine serum and 2% goat serum blocking buffer for 1-2 hours following cryo-sectioning. Another series of PBS washes (3-4 times for 5 minutes each) were performed, and slices were stained with a rabbit polyclonal anti-PCNA primary antibody (Cell Signaling, 2586t) at a concentration of 1:500. All slides were incubated overnight at 4 °C. Following primary antibody incubation, an additional 3-4 PBS washes were performed, and slices were incubated for 2 hours at room temperature in goat anti-rabbit Alexa Fluor-555 secondary antibodies (Thermo Fisher, REF: A21428 Lot: 2633537) diluted at a 1:600 concentration. Secondary antibodies were washed in PBS, then the slides were mounted with one drop of histological mounting medium (Permount SP15-500 toluene solution). Slides were cured in the fridge for 24 hours, then sealed with clear nail polish to protect the tissue from decaying. SOD1 staining followed the same protocol and the primary (GeneTex: GTX637934, lot # 44977) and secondary antibodies applied were at a concentration of 1:1,000.

### Confocal, data acquisition, and analysis

Confocal images were obtained using a Carl Zeiss LSM 800 confocal microscope using a 40x oil-immersion objective lens. Image acquisition settings were standardized across animal groups for all laser channels: Master gain at approximately 560, laser pinhole at approximately 0.40, Digital gain at -3, with 1 μm optical steps, and images were averaged twice. Confocal stacks were analyzed using Imaris software (Oxford Instruments, ver. 9.3) for automated cell counting and colocalization analysis. Cell count of PTN neurons was conducted by selecting the “spots” function, setting the parameters for the region of interest, and using the auto count function. Some cells were manually counted when software errors were identified. The average ratios and standard deviations of each social group (communals, isolates, dominants, and subordinates) were calculated. Volumetric analysis of PTN cell size was conducted using the Imaris digital rendering cell surface function. Figures were compiled in Adobe illustrator. Schematic illustrations were constructed using BioRender.com.

### Statistical analysis

All data were tabulated and organized using Microsoft Excel and GraphPad Prism (v10). Quantifications from confocal images were analyzed using Fiji (ImageJ), and all figures were generated in GraphPad Prism. One-way ANOVA was conducted and followed by Tukey’s Multiple Comparison post-test to compare groups. A statistical threshold of *P* < 0.05 was used for all statistical tests. Data are presented as mean ± standard error of the mean (SEM), and statistical significance is indicated in figure legends with asterisks and corresponding *p*-values. The R statistical environment was used to conduct principal component analysis (PCA) using the prcomp() function (R version 4.3.3, 2024; R Studio version 2023.12.1+402) (R Code Team, 2024). We used packages ggfortify (Tang et al., 2023), factoextra (https://CRAN.R-project.org/package=factoextra), ggplot2 (Wickham, 2016) to visualize the results. The PCA was used to examine and visualize clustering of individuals according to social status groups based on the set physiological and behavioral variables measured. Each sample is represented as a point (or projection) and arrows represent the correlations of each variable to the principal components.

## RESULTS

### Social dominance increases PCNA expression in PTN dopaminergic neurons

The PTN contains three primary DA clusters: the posterior tuberculum anterior rostral (PTar) and posterior tuberculum anterior caudal (PTac) neural clusters project posteriorly into the mesencephalic region to directly modulate the startle escape and swim circuits (Figure 1A-C) (Mu et al., 2012; Ryczko et al., 2013; Haehnel-Taguchi et al., 2018). One distinguishing feature of PTar/PTac neurons is their large almond-shaped somata by comparison with the third neighboring PTp (posterior tuberculum posterior) DA cluster. These morphological features facilitate their identification and differentiation. First, we verified that social dominance modulates the number of PTar/PTac cells of the PTN by analyzing their numbers after 14 days of social interactions. We found that the number of DAT cells was significantly higher in dominant than subordinates and communal zebrafish (Figure 1D; Com n=17, Dom n= 26, Sub n= 24; One-way ANOVA, P=0.0004, One-way ANOVA, Tukey’s Multiple Comparison post-test) consistent with the prior report by Heagy and colleagues (Heagy et al., 2024).

**Figure 1.**
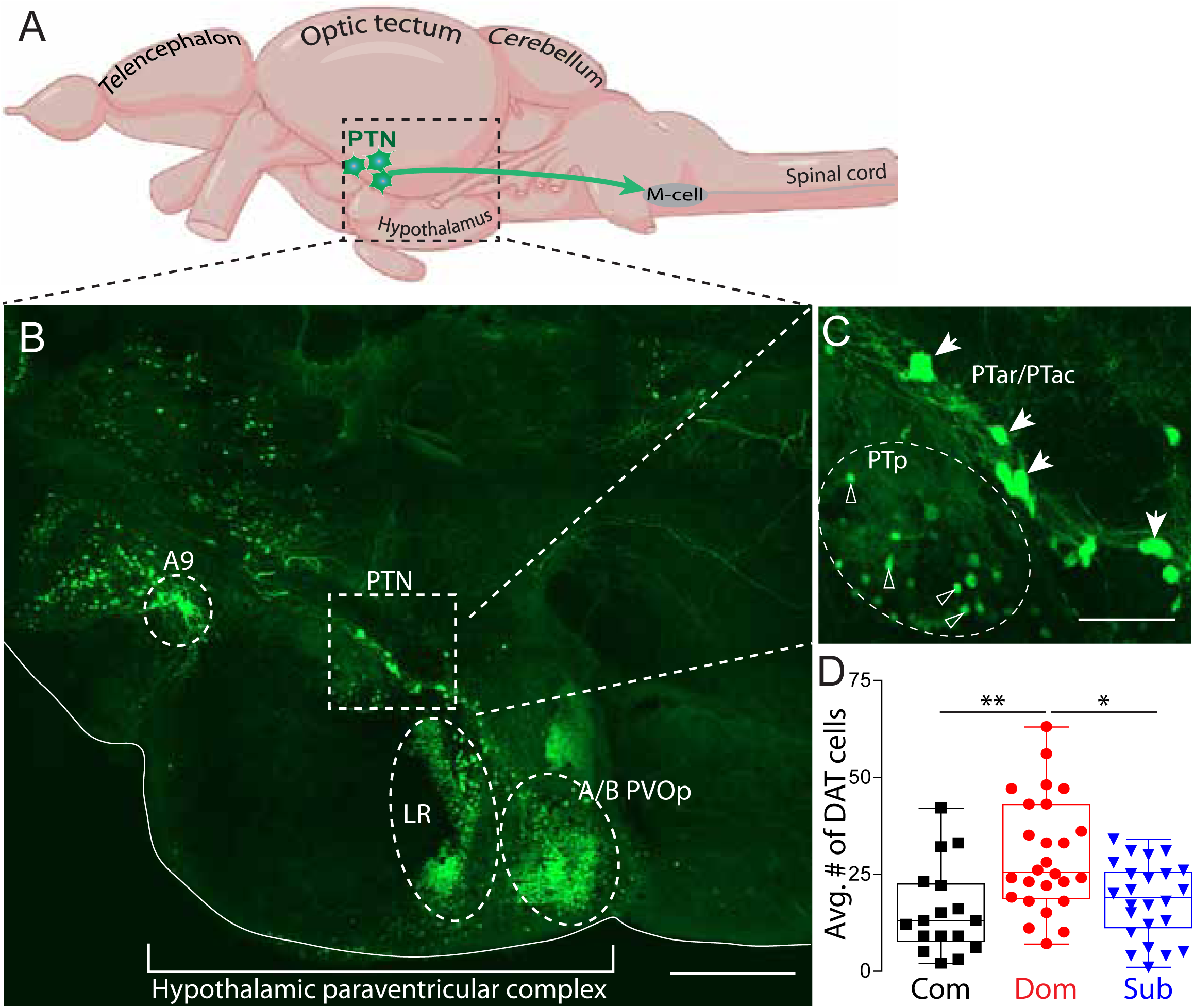
PTN cell number is socially regulated. (A) Schematic sagittal view of zebrafish approximate location of diencephalic posterior tubercular nucleus and projection into spinal cord in relation to the Mauthner neuron (M-cell). (B) a tiled confocal 30 µm maximum projection of the diencephalic and hypothalamic dopamine nuclei including the anterior rostral/anterior caudal PT (PTar/PTac) cluster of the PTN, lateral recess (LR) and paraventricular organ posterior parts A/B (A/B PVOp (Scale 40 µm). (C) Zoomed image of highlighted part of the PTN shown in part B. Filled arrowheads point to the type I PTar/PTac cells. Open arrowheads point to the type II PTp cells (Scale 10 µm). (D) Average number of PTar/PTac DA cells in communal, dominant and subordinate zebrafish (Com n=17, Dom n= 26, Sub n= 24; One-way ANOVA, P=0.0004, One-way ANOVA, Tukey’s Multiple Comparison post-test; \**P*<0.05, \*\**P*<0.01).

Next, we tested whether dominance promotes the addition of new neurons by examining cell proliferation using proliferating cell nuclear antigen (PCNA). PCNA is a DNA polymerase accessory protein expressed during the late G1 and S phases of the cell cycle, making it a reliable marker for identifying actively dividing cells (Zupanc and Horschke, 1995; Grandel and Brand, 2013). In zebrafish, adult cell proliferation occurs in defined proliferative zones throughout the brain (Ming and Song, 2011). Increased PCNA expression in dominant individuals would indicate that social experience enhances cell proliferation in the PTN potentially explaining the elevated number of dopaminergic neurons observed. This approach allows for the spatial localization of proliferating DA cells within the PTN. We found dominant fish exhibited robust PCNA expression that colocalized with PTN neurons (Figure 2A, Movie 1), indicating ongoing cell proliferation. Similarly, communal fish showed moderate PCNA expression, with colocalization in a subset of dopaminergic neurons. However, subordinate fish exhibited less PCNA expression, with minimal overlap between PCNA and PTN neurons. Quantification of PCNA⁺/GFP⁺ cells revealed a significant increase in proliferating dopaminergic neurons in dominant fish compared to subordinate counterparts (Figure 2B; Com n=6, Dom n= 8, Sub n= 8; One-way ANOVA, *P* =0.0029, Tukey’s Multiple Comparison post-test). This result supports the conclusion that social dominance enhances proliferative activity within dopaminergic populations in the PTN.

**Figure 2.**
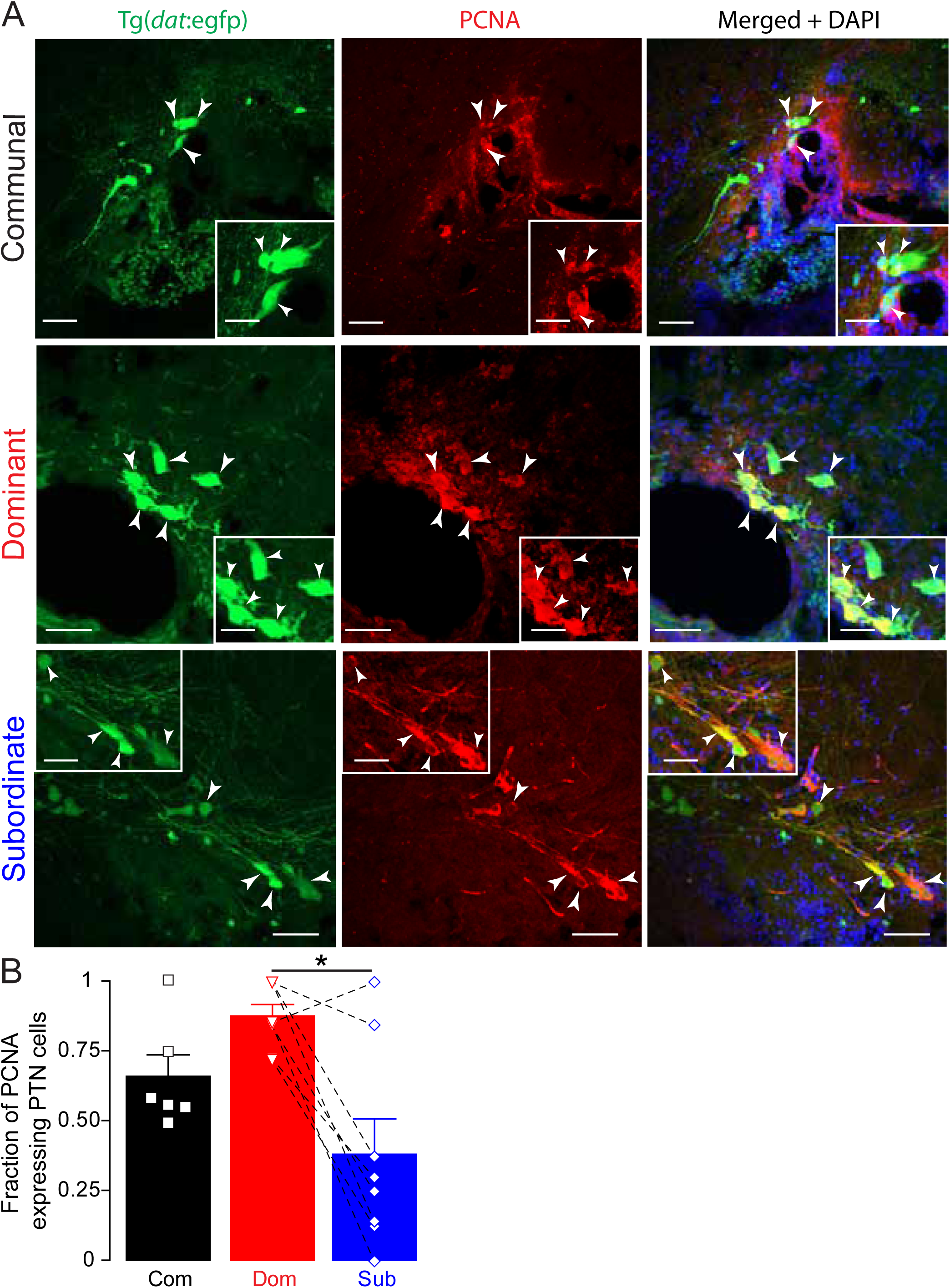
Social dominance affects PCNA expression in PTN neurons. (A) Confocal projections of PCNA expression in a representative communal, dominant, and subordinate Tg(*dat*:egfp) zebrafish. Merged channel also shows DAPI nuclear staining in blue. Arrowheads point to PCNA expressing PTN somata. Scale bar = 20 µm; insets = 5 µm. (B) Summary graph of the fraction of PTN neurons that express PCNA. Symbols represent individual samples; bars represent averages, error bars represent SEM. Dashed lines show pairwise comparison connecting each dominant to its subordinate counterpart. (Com n=6, Dom n= 8, Sub n= 8; One-way ANOVA, *P=0.0029*, One-way ANOVA, Tukey’s Multiple Comparison post-test; \**P*<0.05).

Next, we used bromodeoxyuridine (BrdU) for temporal labeling of cells undergoing mitosis, a robust cell proliferation marker used in several vertebrate models including zebrafish (Ming and Song, 2011; Dunlap et al., 2021). By administering BrdU during defined periods (days 8 & 10 during pairing: the critical time points at which significant differences in cell numbers emerge) and subsequently staining tissue, we were able to identify and quantify cells that are in the S-phase at the time of BrdU incubation (Figure 3A). BrdU complements PCNA by providing a temporal snapshot of cell birth. Together, these two methods provide robust and complementary approaches for cell proliferation and enable assessment of how proliferation correlates with social dominance. We found robust colocalization of BrdU within the PTN DA neurons in dominant zebrafish indicating recent proliferation of dopaminergic neurons (Figure 3B, Movie 2). In contrast, socially communal and isolated zebrafish showed similar BrdU expression (Figure 3B, second row), while subordinate fish exhibited the least amount of BrdU incorporation, with very few colocalized cells (Figure 3B, bottom row).

**Figure 3.**
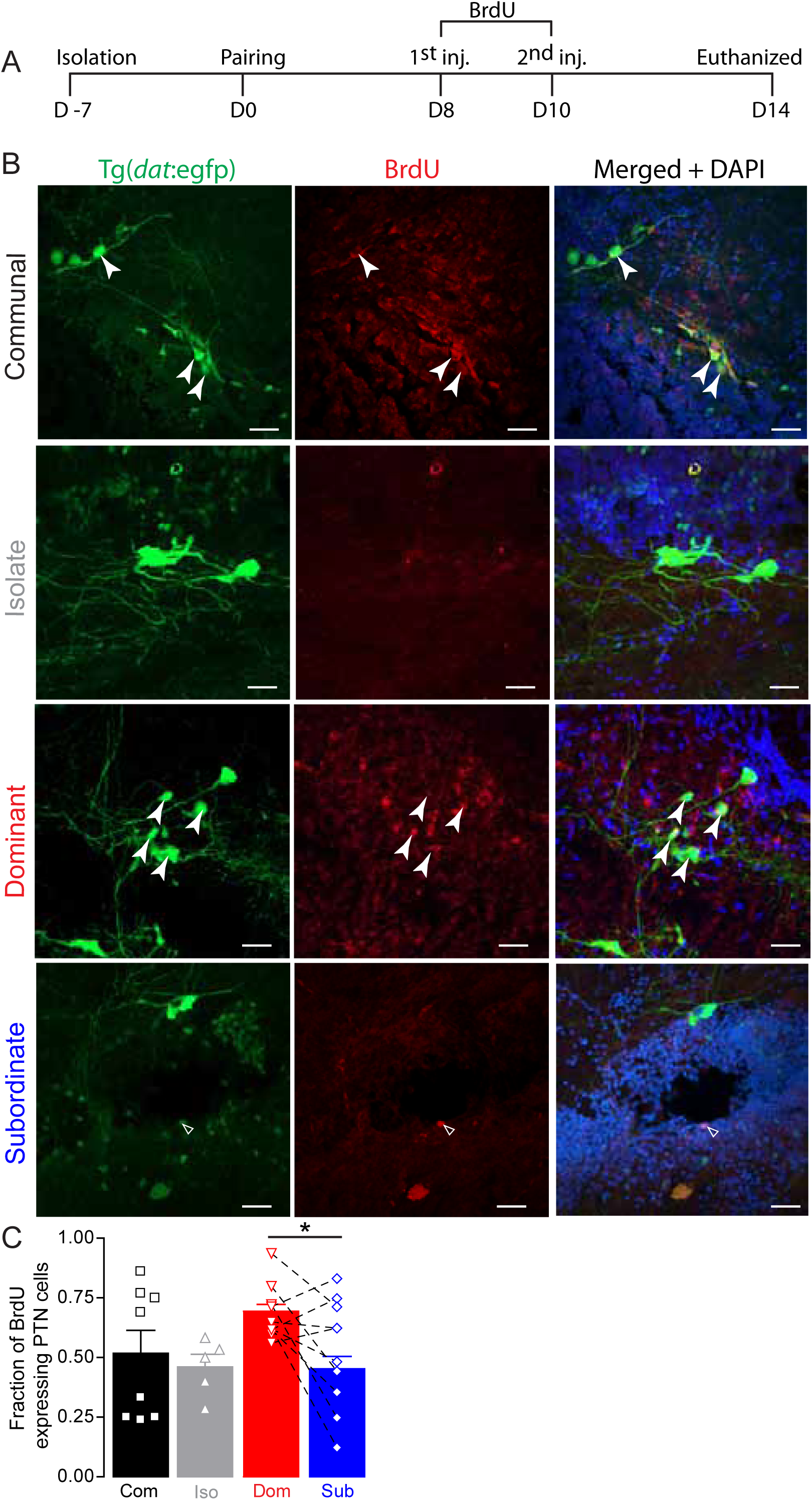
Social dominance affects BrdU expression in PTN. (A) Timeline of behavioral setup, BrdU injections, and tissue extraction. (B) Confocal projections of BrdU expression in a representative communal, isolate, dominant and subordinate Tg(*dat*:egfp) zebrafish. Merged channel also shows DAPI nuclear staining in blue. Closed arrowheads point to BrdU expressing PTN somata; open arrowheads point to none PTN DAT neurons expressing BrdU. Scale bar = 20 µm; insets = 5 µm. (C) Summary graph of the fraction of PTN neurons that express BrdU. Symbols represent individual samples; bars represent averages, error bars represent SEM. Dashed lines show pairwise comparison connecting each dominant to its subordinate counterpart. (Com n=8, Iso n = 5, Dom n= 10, Sub n= 10; One-way ANOVA, Tukey’s Multiple Comparison post-test *P=0.0473*; \**P*<0.05).

Quantitative analysis of BrdU expression within the PTN DA cells across groups revealed a significant effect of social condition on cell proliferation (Figure 3C). Dominant animals exhibited the highest expression level within the PTN neurons, with communal fish showing a moderate BrdU level. Isolated and subordinate animals showed fewer BrdU expression, but statistically the levels of expression were not significant from communal fish. However, there were statistical differences in BrdU expression among the groups and between dominant and subordinate zebrafish (Com n=8, Iso n = 5, Dom n= 10, Sub n= 10; One-way ANOVA, *P=0.0473*, Tukey’s Multiple Comparison post-test). Taken together, these results indicate that the dopaminergic system in the PTN exhibits structural plasticity in response to social context. Social dominance appears to selectively promote the generation of new dopaminergic neurons, while subordination and isolation are associated with reduced levels of cellular proliferation.

### Social status influences neurotransmitter identity in PTN dopaminergic neurons

An alternative mechanism that may explain the differences in the number of DA PTN cells may include a switch in neurotransmitter phenotype. Prior studies in mice demonstrated that diencephalic dopaminergic neurons can co-express and switch between dopaminergic and glutamatergic phenotypes (Kawano et al., 2006). Similarly, previous findings in zebrafish showed that PTN neurons co-express dopamine and glutamate (Filippi et al., 2014; Altbürger et al., 2024). However, whether such shifts in neurotransmitter identity occur and whether they are socially regulated, was unclear.

To explore whether social dominance induces a switch in neurotransmitter expression, we used a double transgenic zebrafish line, Tg(*dat*:egfp), Tg(*vglut2a:*rfp) and compared the proportions of glutaminergic and dopaminergic PTN neurons among social groups. A decrease in glutamate (RFP) and an increase in dopamine (EGFP) within the same population suggests that social experience may drive a shift toward a dopaminergic phenotypic identity and vice versa. This type of plasticity may help explain the increase in dopaminergic neurons observed in dominants and could reflect the brain’s adaptation of neural function to support different behavioral roles.

Confocal microscopy showed that a subset of the PTN dopaminergic neurons co-express Vglut2a and Dat in all social phenotypes (Figure 4A). Interestingly, we found that the ratio of Vglut2a expressing cells to DAT cells in subordinate fish was significantly higher compared to dominants and communals but did not differ significantly from those of isolated fish (Figure 4B, Movie 3; Com n=7, Iso n = 4, Dom n= 14, Sub n= 10; One-way ANOVA, *P=0.0174;* Tukey’s Multiple Comparison Test). These results suggest that social experience influences the pattern of neuromodulator expression of the PTN whereby social stress (either isolation or submission) leads to an increase in the ratio of Vglut2a/DAT expressing cells, the functional consequence of which on motivational and motor behavior remains unknown.

**Figure 4.**
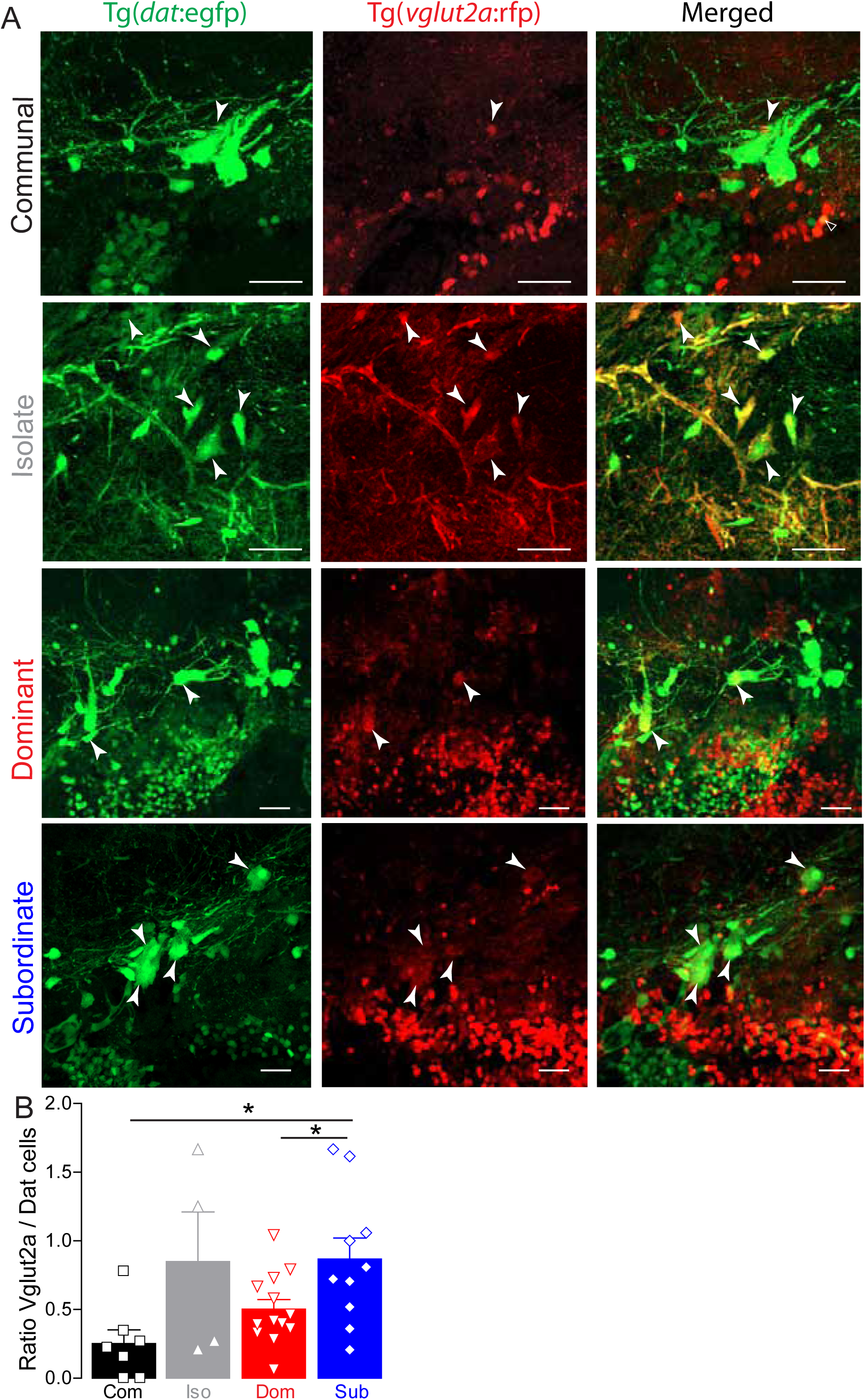
Social dominance shifts PTN neurotransmitter expression pattern. (A) Confocal projections of PTN cells that co-express DAT and VGLUT2A from a communal, isolated, dominant and subordinate double transgenic Tg(*dat*:egfp), Tg(*vglut2a:*rfp) zebrafish. Arrowheads point to PTN cell that co-express DAT and VGLUT2A. Scale bar = 20 µm; insets = 5 µm. (B) Summary graph of the ratio of VGLUT2A expressing cells to those that express DAT. Symbols represent individual samples; bars represent averages, error bars represent SEM. (Com n=7, Iso n = 4, Dom n= 14, Sub n= 10; One-way ANOVA, *P=0.0174*, Tukey’s Multiple Comparison post-test; \**P*<0.05).

### Social Subordination and isolations influence oxidative stress levels

Next, we tested the alternative hypothesis that status-dependent differences in PTN DAT cell number may also be a result of cellular stress that leads to neurodegeneration in subordinates. In many social species, social subordination is often stressful, which may induce oxidative cellular damage and eventual cellular loss. However, whether oxidative stress could contribute to cell loss seen in chronically stressed animals was unknown.

Reactive oxygen species (ROS), generated under metabolic stress, can cause damage to cellular proteins, membranes, and DNA. To counter this, cells express antioxidant enzymes like superoxide dismutase 1 (SOD1), a mitochondrial antioxidant enzyme involved in caspase-independent cell death, which converts superoxide radicals into less harmful molecules (Massaad and Klann, 2011). We hypothesized that elevated SOD1 expression would indicate an attempt to compensate for increased oxidative burden, suggesting that stress-induced neurotoxicity may underlie reductions in neuronal viability. If so, then the observed difference in cell number could be a result of reduced viability and loss of PTN neurons in subordinates because of stress. Moreover, social isolation is stressful for many social species, and prolonged social isolation may also compromise PTN health (Heagy et al., 2024).

To test this hypothesis, we examined expression of SOD1 and quantified its immunoreactivity in Tg(*dat*:egfp) fish across social groups to determine whether stress exposure associated with status or isolation increases oxidative state in dopaminergic neurons. We found that in communal and dominant zebrafish, SOD1 labeling was relatively low, with sparse expression observed in the PTN dopaminergic neurons (Figure 5A, Movie 4). In contrast, isolated fish exhibited robust SOD1 expression, with strong colocalization in many PTN neurons suggesting elevated oxidative stress under socially deprived conditions (Figure 5A). Subordinate fish similarly displayed high SOD1 expression, with many PTN neurons co-labeled for SOD1 (Figure 5).

**Figure 5.**
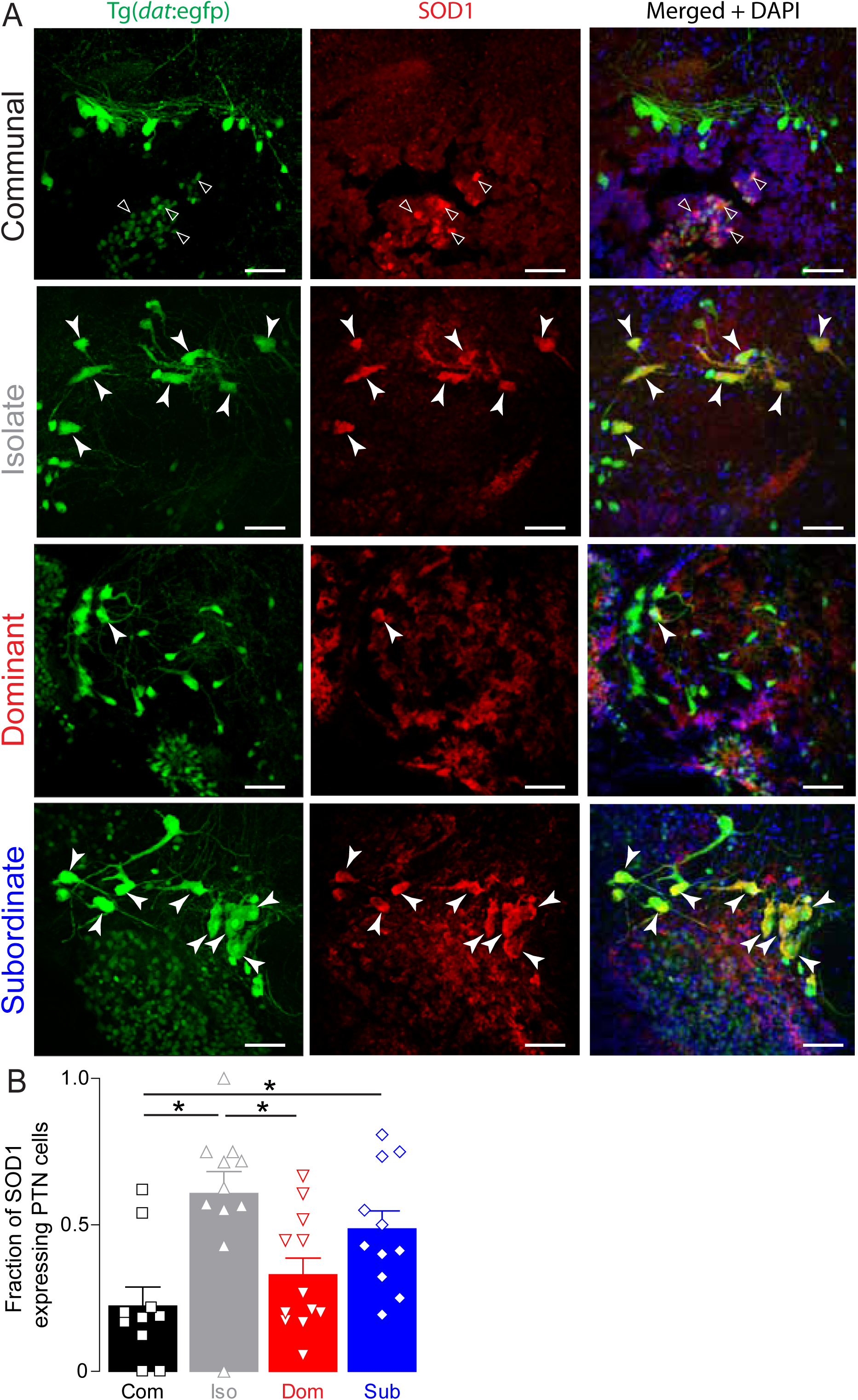
Social isolation and subordination induce oxidative stress in PTN neurons. (A) Representative confocal projections of SOD1 expression in PTN cells of a representative communal, isolated, dominant, and subordinate Tg(*dat:*egfp) zebrafish. Merged channel also shows DAPI nuclear staining in blue. Closed arrowheads point to SOD1 expressing PTN somata; open arrowheads point to none PTN DAT neurons expressing SOD1. Scale bar = 20 µm; insets = 5 µm. (B) Summary graph of the fraction of PTN neurons that express SOD1. Symbols represent individual samples; bars represent averages, error bars represent SEM (Com n=10, Iso n= 11, Dom n= 12, Sub n= 11; One-way ANOVA, *P=0.001*, Tukey’s Multiple Comparison post-test; \**P*<0.05).

Quantification of SOD1 expression revealed that isolated fish exhibited the highest SOD1 expression in PTN neurons, followed by subordinate fish (Figure 5B; Com n=10, Iso n= 11, Dom n= 12, Sub n= 11; One-way ANOVA, *P=0.001*, Tukey’s Multiple Comparison post-test). Intriguingly, dominant and communal animals showed significantly lower SOD1 expression, suggesting a protective or adaptive regulation of oxidative stress in dominants (Figure 5B). These results indicate that social experience impacts cellular oxidative stress in the PTN. Social isolation and subordination are associated with elevated SOD1 expression in dopaminergic neurons, while dominant status appears to be protective. These findings support the hypothesis that social hierarchy modulates cellular stress responses in neuromodulatory systems.

### Multivariate correlative analysis of physiological adaptations to social conditions

To examine how physiological differences correlate with social status, we conducted a multivariate principal component analysis (PCA) to determine main physiological parameters measured that correlate with status (Figure 6). Given the absence of PCNA data in socially isolated zebrafish, we first conducted PCA that included communals, dominants, and subordinates only along with all the physiological parameters measured (BrdU, PCNA, vglut/dat ratio expression, and SOD1) (Figure 6A). The first two principal components explained about 65% of the variance (Figure 6B). The dominant individuals clustered together where the variables PTN cell number, BrdU and PCNA were positively correlated and loaded strongly to PC1 axis. In contrast, the subordinate zebrafish clustered together, where the variable SOD1 expression was positively correlated and strongly loaded to PC2 axis while vglut/dat ratio was negatively loaded to the PC1 axis. Next, we conducted a separate PCA in which we included the social isolates but excluded the PCNA data to determine the main physiological parameters that correlate with status (Figure 6C). The first two principal components explained about 65% of the variance (Figure 6D). The dominant individuals clustered together where the variables PTN cell number and BrdU were positively correlated and loaded strongly to PC1 axis. In contrast, the socially isolated zebrafish clustered together, where the variable SOD1 and cell identity ratio were positively correlated and loaded to PC1 and PC2 axes, respectively.

**Figure 6.**
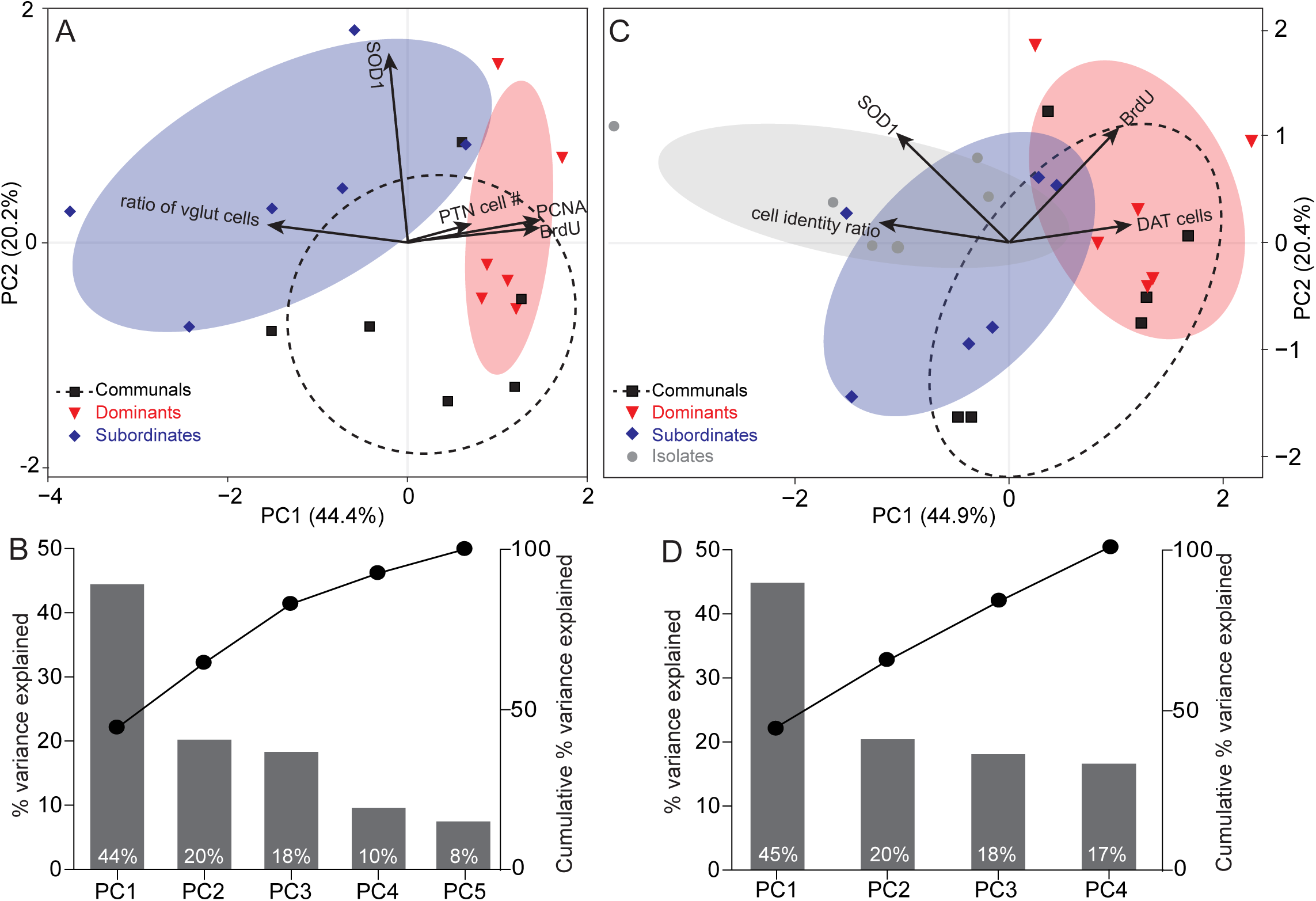
Ordination based on a PCA depicting PTN cell number, BrdU, PCNA, Vglut2a, and SOD1 expression according to social rank. (A) PC1 and PC2 are the first and the second principal components excluding social isolates. (B) Eigenvalues of the principal components excluding social isolates showing the percentage of variation explained by each component. Line plot illustrates the cumulative explained variance. (C) PC1 and PC2 are the first and the second Principal components excluding PCNA as a component. (D) Eigenvalues of the principal components excluding PNCA as a component showing the percentage of variation explained by each component. Line plot illustrates the cumulative explained variance. In A and C, points represent individual samples (n= 6 per social condition), different colors and symbols represent different groups. Ellipses represent 68% confidence intervals of core regions. Loading vectors represent original variables, the directions of arrows represent correlation between original variable and principal components, lengths represent association of original data to principal components. In B and D components are listed in descending order (highest to the lowest).

Collectively, the multivariate analysis corroborates our observations by revealing distinct clusters among the social groups emphasizing the divergence in physiological and behavioral activities as animals ascend or descend in social rank or experience prolonged periods of isolation. Social dominance promotes dopaminergic neuronal growth and proliferation (BrdU and PCNA); while social subordination or isolation are detrimental and induce cellular stress (SOD1) and a shift in neurotransmitter identity.

## DISCUSSION

The primary objective of our study was to examine the cellular mechanisms underlying status-dependent plasticity in PTN cellular number. Our results demonstrate that social status reconfigures both the structural organization and functional activity of the PTN, offering a cellular foundation for how social experience influences behavior. Specifically, our results suggest a model where dominant zebrafish exhibit a higher number of PTN dopaminergic neurons, driven by increased levels of proliferation and changes in neuromodulator expression (Figure 7). Conversely, the PTN of subordinate and socially isolated zebrafish display reduced cell proliferation and enhanced levels of oxidative stress, suggesting that stress-induced mechanisms may reduce neuronal viability and contribute to the decline in neuronal number (Figure 7).

**Figure 7.**
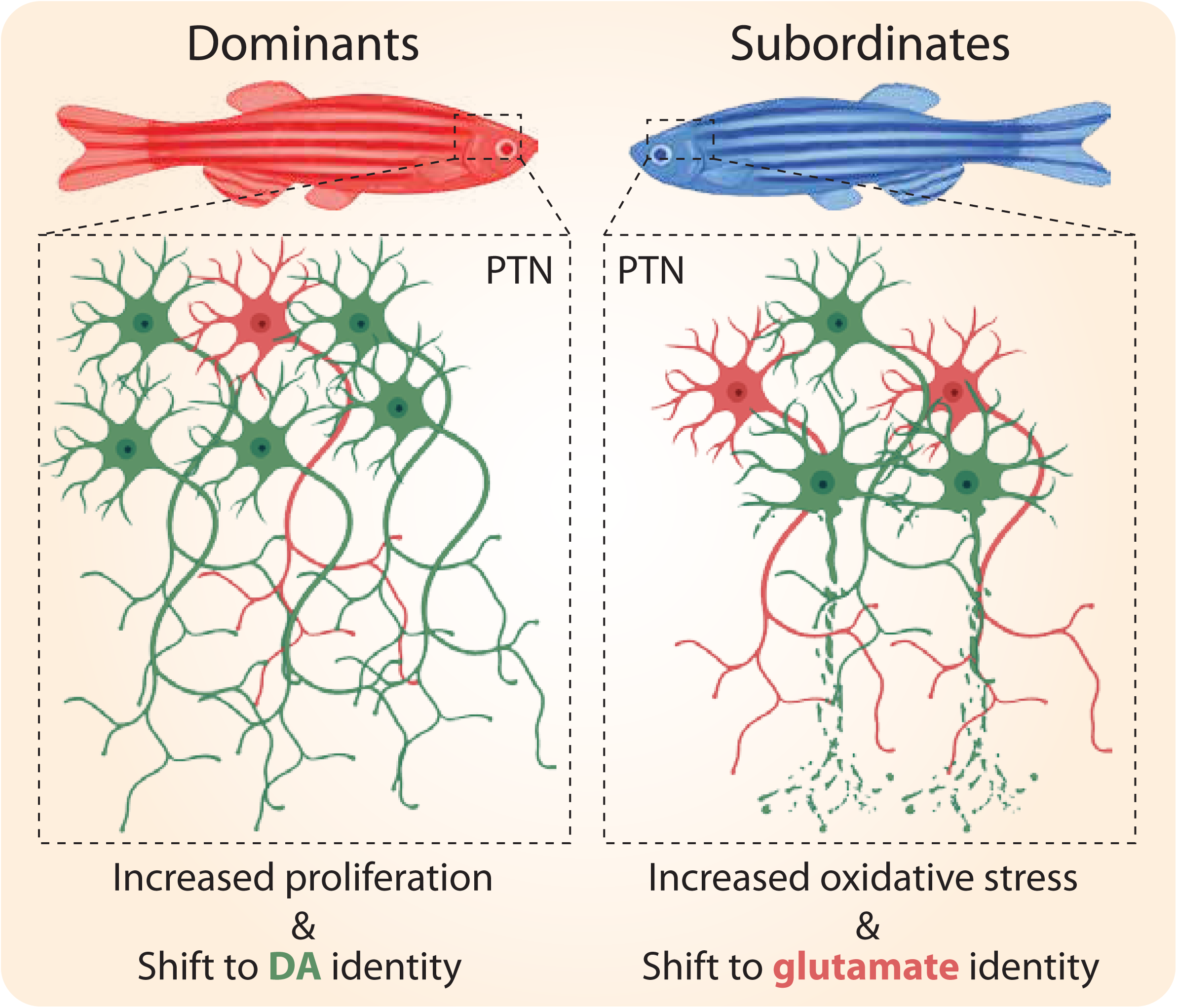
Schematic illustration of the proposed model of how social dominance affects PTN morphological architecture. Social dominance promotes cell proliferation and proliferation of dopaminergic neurons; simultaneously, existing PTN neurons shift their neurotransmitter identity from glutaminergic to dopaminergic. Collectively, this is likely to increase DA release to modulate postsynaptic targets. Social stress due to subordination or isolation induces oxidative stress and reduction in PTN neuronal viability; simultaneously, existing PTN neurons shift their neurotransmitter identity from dopaminergic to glutaminergic expression. Collectively, this is likely to reduce DA release and lead to differences in modulatory signaling of postsynaptic targets compared to dominants.

### Social dominance and the regulation of cell proliferation

A central finding of our work is that social rank directly modulates the birth of new dopaminergic neurons in the PTN. Dominant zebrafish gradually accumulate more PTN neurons and show enhanced proliferation, as evidenced by significantly higher PCNA and BrdU incorporation. This result aligns with the broader findings that social environments act as potent drivers of structural brain plasticity. In African cichlid fish, social ascent triggers rapid changes in gonadotropin-releasing hormone (GnRH) neuron size and physiological profile, demonstrating that the social environment can fundamentally remodel neuromodulatory systems to support status-specific behaviors (Hofmann, 2003; Maruska and Fernald, 2011).

Our results extend this concept to the dopaminergic system, suggesting that the socially plastic brain utilizes adult cell proliferation to scale its modulatory capacity in response to increased social demand. These differences, which emerge around 7–14 days post-dominance formation, suggest that PTN neuronal number is not predetermined but is shaped by extended social exposure. This is consistent with prior work in our lab showing that dopamine cell number in diencephalon is socially regulated: socially dominant zebrafish have more dopamine neurons in specific descending clusters than subordinates (Heagy et al., 2024).

The reduction of cell proliferation in subordinate and isolated zebrafish further highlights the sensitivity of these proliferative zones to social stress. In rodents, chronic social defeat is well-known to suppress hippocampal cell proliferation through the activation of the hypothalamic-pituitary-adrenal (HPA) axis and the subsequent release of glucocorticoids (Czéh et al., 2001). Our findings suggest a similar mechanism could be at play in the zebrafish PTN, where social stress reduces neurogenesis and prevents the expansion of the dopaminergic population.

A critical consideration in interpreting the social remodeling of the PTN is the high capacity for constitutive and regenerative adult neurogenesis in zebrafish, a trait that distinguishes teleost fish from most mammalian lineages (Lindsey and Tropepe, 2006; Ganz and Brand, 2016). In zebrafish, neural stem cell niches are distributed throughout the brain, including the posterior hypothalamus and periventricular zones adjacent to the PTN, allowing for the continuous integration of new neurons into functional adult circuits (Grandel et al., 2006). Our observation that social dominance significantly upregulates cell proliferation and the recruitment of new dopaminergic neurons suggests that social status acts as a potent regulator, similar to how environmental enrichment influences other teleost brain regions (von Krogh et al., 2010). However, because adult neurogenesis is largely restricted to the subventricular and subgranular zones in mammals (Bond et al., 2015), the extensive structural turnover we observed may represent a mechanism of plasticity that is particularly pronounced in teleosts. While the underlying drivers of this remodeling such as social stress or competitive success are likely conserved across vertebrates, the specific cellular execution in zebrafish appears multifaceted and relies on the addition of neurons rather than purely synaptic or biochemical modifications.

### Phenotypic plasticity: Neurotransmitter switching

While cell proliferation explains the addition of neurons, our observation of status-dependent shifts in VGLUT2a and DAT expression ratios suggests that the PTN also employs neurotransmitter switching—a more rapid form of plasticity. Subordinate and socially isolated zebrafish displayed a significantly higher ratio of VGLUT2a expression in DAT cells compared to dominants. This suggests that social stress may favor a glutamatergic or co-expressing phenotype the functional consequence of which remains unresolved. One possibility is the social stress vs. trophic reward framework gleaned from our PCA suggesting that dominance is not just the absence of stress, but a positive trophic state. The tight correlation among DAT cell number, PCNA and BrdU on the PCA loading plot suggests that dominance-induced neurogenesis is specifically geared toward expanding the dopaminergic population. This supports a hypothesis where social success reinforces the PTN-to-striatum reward circuitry via continuous cellular addition. Conversely, the PCA highlights a reciprocal relationship between dopaminergic and glutaminergic identity suggesting a potential phenotypic switch. In subordinates, the chronic social stress evidenced by the SOD1 loading may trigger a survival-based plastic response where neurons downregulate dopamine, which is highly metabolically active and oxidative-prone and upregulate glutamate to maintain basic circuit stability while minimizing further oxidative damage (Ni and Ernst, 2022; Flores-Ponce and Velasco, 2024). It is noteworthy that isolated zebrafish cluster closer to subordinates than to dominants on the PCA suggesting that social isolation and social defeat share a common oxidative neurobiological signature, distinct from the unique trophic signature of dominants.

In mice, diencephalic dopaminergic neurons have been shown to switch their neurotransmitter identity in response to photoperiod changes or environmental stimuli (Dulcis and Spitzer, 2008). Such switching allows an animal to reconfigure circuit output without the metabolic cost or time delay associated with generating new cells. Thus, structural plasticity in the PTN likely arises through both proliferation and neurotransmitter switching. However, more subtle shifts, such as partial co-expression or modulation of release machinery, may occur below the detection threshold with significant functional consequences on behavior (Heagy et al., 2024). The increase in glutamatergic signaling in subordinates and isolates might represent an adaptive shift in circuit gain, perhaps modulating locomotor drive or startle sensitivity to better suit a submissive and stressed behavioral strategy (Clements et al., 2023). Therefore, while the PTN acts as a conserved hub for translating social experience into behavioral drive, the high degree of cellular turnover we observed emphasizes that zebrafish may utilize multifaceted mechanisms to achieve circuit state; while mammalian systems are maintained through more localized synaptic plasticity. Recognizing this distinction is essential for modeling how social status-induced changes in dopamine signaling translate across diverse vertebrate architectures.

### Oxidative stress and neuronal resilience

The elevated SOD1 expression in isolated and subordinate zebrafish suggests that social rank influences the cellular oxidative state. Social isolation, particularly, appeared to be the most metabolically taxing condition. In mammals, social isolation is a recognized stressor that increases reactive oxygen species (ROS) and leads to dopaminergic dysregulation, particularly in the prefrontal cortex and striatum (Möller et al., 2011; Liu et al., 2012; Schiavone et al., 2013; Shao et al., 2015; Lampert et al., 2017; Mumtaz et al., 2018). The robust SOD1 expression in subordinates and isolates may reflect a compensatory attempt to detoxify superoxide radicals. However, if this oxidative burden is prolonged and exceeds cellular capacity, it may lead to the degeneration or apoptosis of PTN neurons. This provides a potential explanation for why subordinates and isolates do not simply maintain a static population of neurons but rather show a decline in cell number over time. It is possible that neuronal death and suppressed cell proliferation act simultaneously; subordinates may produce fewer neurons while simultaneously exposing existing ones to oxidative stress that leads to a net decline in PTN neuronal number. Future work could apply markers of apoptosis (e.g., caspase-3, TUNEL) to test this hypothesis. Dominant status, conversely, appears to be neuroprotective, with low SOD1 levels suggesting a more stable cellular environment that supports neuronal survival and high activity levels.

### Functional integration and behavioral scaling

The multivariate PCA revealed a clear divergence in physiological profiles based on social rank. The correlation of PTN cell number and BrdU expression in dominants—combined with PCNA expression—indicates that these neurons are functionally engaged. Moreover, Heagy and colleagues have shown that PTN cells in dominants show increased PS6 expression indicative of a higher cellular activity compared to subordinates and isolated zebrafish reflecting sustained cellular engagement (Heagy et al., 2024). Collectively, the results that dominant zebrafish display elevated PTN activity indicate that they not only generate more dopaminergic neurons but also activate them more robustly. Conversely, PTN cellular number and PS6 activity is reduced in stressed fish correlating strongly with the elevated SOD1 expression suggesting that elevated oxidative stress may play a detrimental role in reducing PTN viability.

In zebrafish, the PTN is implicated in sensorimotor integration, particularly in the modulation of the Mauthner cell mediated startle response. Social status shifts behavior: dominants increase swimming and suppress startle sensitivity, while subordinates reduce swimming and heighten startle responsiveness. These differences are regulated by synergistic interactions among dopaminergic, glycinergic, and GABAergic inputs that converge on the Mauthner circuit (Clements et al., 2023). Our results suggest that the PTN serves as an upstream supply-side regulator. Dominants have more active PTN neurons, providing stronger dopaminergic drive onto the motor circuit. This likely feeds into the drd1b molecular switch in downstream glycinergic neurons, enabling stronger inhibition of the M-cell and suppressing startle responses (Clements et al., 2023). In subordinates, having fewer and less active PTN cells would reduce dopaminergic signaling, disinhibiting the M-cell and increasing startle sensitivity. This model supports the idea that upstream anatomical plasticity coupled with downstream neuromodulatory changes to route behavioral outputs as an adaptation to social rank.

In conclusion, our results show that the PTN is a highly dynamic neural hub that adapts its structure and function to the social environment. By integrating cell proliferation, neurotransmitter switching, and stress-response molecular mechanisms, the PTN enables zebrafish to calibrate their behavioral output to their social standing.

## Acknowledgements

This study was funded by the NIH (grant no. R15MH141535) and NSF (grant no. 1754513). We thank Drs Karen Litwa and Lauren Anllo for constructive feedback; Dr Timothy Erickson for providing the Tg(*dat*:egfp) line; Ms. Feliciti Gunter for technical assistance, and the Imaging Core Facility at ECU Biology department for instrument access.

